# Sex-specific Genetic Regulatory Effects in Chickens

**DOI:** 10.64898/2026.02.26.708179

**Authors:** Haihan Zhang, Bingjin Lin, Yang Xi, Di Zhu, Chen Peng, Junhao Tu, Huichao Liu, Dailu Guan, Qingyuan Ouyang, Bogong Liu, Chenyu Fan, Zichao Song, Xing Meng, Houcheng Li, Bingxing An, Yanan Wang, Jingsheng Lu, Mingshan Wang, Xuemei Lu, Jinyan Teng, Mian Gong, Leilei Cui, Bingjie Li, Dominic Wright, Marta Gòdia, Ole Madsen, Ruidong Xiang, Yang I. Li, Iuliana Ionita-Laza, Yulong Yin, Xiao Wang, Dan Hao, Qin Zhang, Weijie Zheng, Xiaoning Zhu, Songchang Guo, Zehe Song, Kun Xie, Guoqiang Yi, Zhixiu Wang, Zhuanjian Li, Chenghao Zhou, Yuzhe Wang, Congjiao Sun, Corie Darrington, Shu-Hua Hsu, Ning Yang, Xiaoxiang Hu, Huaijun Zhou, Xin Zhao, Monique G.P. van der Wijst, Jian Zeng, Lingbin Liu, Shourong Shi, Yali Hou, Pengju Zhao, Guiping Zhao, Lingzhao Fang, Xi He

**Affiliations:** College of Animal Science and Technology, Hunan Agricultural University, Changsha 410128, China; Hainan Institute, Zhejiang University, Yongyou Industry Park, Yazhou Bay Sci-Tech City, Sanya 572000, China; Center for Quantitative Genetics and Genomics (QGG), Aarhus University, Aarhus, 8000, Denmark; College of Life Science, Sichuan Agricultural University, Ya’an 625014, China; State Key Laboratory of Livestock and Poultry Breeding, Guangdong Provincial Key Lab of Agro-Animal Genomics and Molecular Breeding, College of Animal Science, South China Agricultural University, Guangzhou 510642, China; State Key Laboratory of Animal Biotech Breeding, Institute of Animal Sciences, Chinese Academy of Agricultural Sciences (CAAS), Beijing, China; Yuelushan Laboratory, Changsha, 410128, China; Poultry Institute, Chinese Academy of Agricultural Sciences, Yangzhou, Jiangsu 225125, China; Kunming Institute of Zoology, Chinese Academy of Sciences, Kunming, Yunnan 650223, China; Metabolic Control and Aging, Human Aging Research Institute (HARI) and School of Life Science, Nan-chang University, Jiangxi Key Laboratory of Human Aging and Jiangxi Province Key Laboratory of Aging and Disease, Nanchang 330031, China; School of Life Sciences, Nanchang University, Nanchang, China; Department of Animal Biosciences, Sveriges Landbruks Universitet (SLU) Ultuna, Uppsala 75007, Sweden; Animal Breeding and Genomics, Wageningen University and Research, Wageningen, 6700 AH The Netherlands; School of Applied Systems Biology, La Trobe University, Bundoora, VIC 3083, Australia; Section of Genetic Medicine, University of Chicago, Chicago, IL, USA; Department of Human Genetics, University of Chicago, Chicago, IL, USA; Department of Biostatistics, Columbia University, New York, NY, USA; Laboratory of Animal Nutritional Physiology and Metabolic Process, Key Laboratory of Agroecological Processes in Subtropical Region, National Engineering Laboratory for Pollution Control and Waste Utilization in Livestock and Poultry Production, Institute of Subtropical Agriculture, Chinese Academy of Sciences, Changsha, 410125, China; Institute of Animal Science and Veterinary Medicine, Shandong Academy of Agricultural Sciences, Jinan, 250100, China; Poultry Institute, Shandong Academy of Agricultural Sciences, Ji’nan 250100, China; College of Animal Science and Technology, Shandong Agricultural University, Taian, Shandong 271018, China; State Key Laboratory of Genome and Multi-omics Technologies, Shenzhen Branch, Guangdong Laboratory of Lingnan Modern Agriculture, Key Laboratory of Livestock and Poultry Multi-omics of MARA, Agricultural Genomics Institute at Shenzhen, Chinese Academy of Agricultural Sciences, Shenzhen, China; College of Animal Science and Technology, Southwest University, Chongqing 400715, China; Key Laboratory for Animal Genetics & Molecular Breeding of Jiangsu Province, College of Animal Science and Technology, Yangzhou University, Yangzhou 225009, China; College of Animal Science and Technology, Henan Agricultural University, Zhengzhou 450046, China; National Chickens Genetic Resources, Jiangsu Institute of Poultry Science, Yangzhou 225125, China; College of Animal Science and Technology, China Agricultural University, Beijing 100193, China; State Key Laboratory of Animal Biotech Breeding, College of Biological Sciences, China Agricultural University, Beijing 100193, China; Department of Animal Science, University of California, Davis, CA, 95616, USA; Department of Animal Science, McGill University, Quebec, H9X3V9, Canada; Department of Genetics, University of Groningen, University Medical Center Groningen, Groningen, the Netherlands; Institute for Molecular Bioscience, The University of Queensland, Brisbane, QLD 4072, Australia

## Abstract

Sexual dimorphism is a defining vertebrate feature, yet its sex-specific molecular architecture remains poorly understood. Here we established a sex-balanced, uniformly reared chicken cohort to map this landscape, integrating individual whole-genome sequencing with 7,969 bulk and 779,380 single nucleus transcriptomes across 32 tissues from 280 birds. We identified 495,098 independent expression quantitative trait loci for 20,194 genes, including 10,937 loci modulated by cell-type composition. Notably, 340 genes were regulated by 449 loci in a sex-dependent manner, significantly enrichment in endocrine tissues like adipose and the adrenal gland. Furthermore, we fine-mapped 1,219 structural variants, demonstrating their unique roles to tissue- and sex-specific expression beyond SNPs. Ultimately, we showed the utility of these regulatory effects in elucidating the molecular basis of metabolism and complex traits in both chickens and humans. This comprehensive atlas of regulatory effects provides profound insights into the genomic and molecular basis of sexual dimorphism in vertebrates.

## Main Text

Sexual dimorphism in morphology, physiology, and behavior is a pervasive hallmark of vertebrate evolution, including humans (*1, 2*). These phenotypic divergences are fundamentally anchored in sex-biased genetic architectures that dictate disease susceptibility and evolutionary fitness. In humans, sex-stratified genome-wide association studies (GWAS) have uncovered numerous loci with dimorphic effects on complex traits and diseases, exemplified by the disproportionate genetic risk for major depressive disorder in females (*3–6*). Despite the profound clinical and biological implications of these sex-biased effects, the underlying molecular mechanisms remain elusive. The domestic chicken (*Gallus gallus domesticus*) provides a powerful model to bridge this gap. Centuries of intensive selection for sex-specific complex traits of economic importance—such as egg production in females and meat yield in males—have magnified sexual dimorphism, creating a stark phenotypic contrast that is ideal for genetic dissection (*9–11*). Beyond its critical role in global food security, the chicken serves as a cornerstone of avian biology and a well-established surrogate for human biomedical research, particularly in development and immunology (*7, 8*). Importantly, birds utilize a ZW sex-determination system that provides an evolutionary contrast to the mammalian XY system. Unlike mammals, chickens lack the chromosome-wide meiotic inactivation, leading to pronounced Z-chromosome transcriptomic disparities and incomplete dosage compensation (*12–15*). This divergence offers an unparalleled lens to investigate how the genome manages sex-specific regulation across diverse phenotypes (*16–19*).

Current efforts to map the sex-specific regulatory landscape have primarily focused on humans and a few model organisms (*20–22*). For instance, the human Genotype-Tissue Expression (GTEx) project identified sex-biased regulatory variants across 44 human tissues, providing critical insights into female-specific risk loci for diseases like breast cancer (*22*). However, human studies are intrinsically limited; significant sex-specific effects are often confounded by lifestyle, diet, and environmental heterogeneity (*22*). In chickens, while sex-interaction effects have been reported for critical phenotypes like feed efficiency and Marek’s disease susceptibility (*23, 24*), a comprehensive regulatory atlas has not yet been established. Although the pilot phase of the ChickenGTEx project explored tissue-level regulation based on publicly available data, it lacked the statistical power and controlled experimental design necessary to disentangle sex-biased regulation from confounding factors such as projects, breeds and developmental stages (*25*). Consequently, the extent to which genetic variation interacts with sex to drive the molecular architecture of complex traits in birds and how this mirrors or differs from mammalian paradigms remain fundamental open questions in evolutionary and agricultural genomics.

To address these limitations, here we established the ChickenSexGTEx resource (**Fig. 1**), where we utilized a genetically diverse population of local chickens, that was maintained for eight generations of random mating. From this cohort, 1,398 birds were reared under strictly uniform environmental conditions to eliminate external noise. We systematically sampled 32 primary tissues from a sex-balanced sub-cohort of 280 individuals (150 males and 130 females) at the age of 90 days, generating 8,265 bulk RNA-seq datasets paired with individual whole-genome sequencing (WGS, 30×) and rich metadata. including body weight, feed intake, body composition and nutrient digestibility metrics. To ensure physiological fidelity, all tissues were harvested within 20 minutes of slaughter to capture near-native transcriptomic profiles. To refine the cellular resolution of this atlas, we integrated 93 single-nucleus RNA-seq (snRNA-seq, 779,380 nuclei) datasets, enabling the computational deconvolution of cell-type-specific signals across all tissues. This multi-layered framework allowed us to identify regulatory variants—including both SNPs and structural variants (SVs)—that exhibit sex-, tissue-, and cell-type-dependent effects. By integrating this regulatory map with 865 serum lipid metabolites and 26 complex traits in 1,575 chickens of the same population, we elucidated the molecular logic by which sex-specific variants drive higher-order phenotypes. Finally, we demonstrated the evolutionary conservation of regulatory effects between chickens and humans, and the potential of ChickenSexGTEx resources in illustrating human complex traits. This work provides an invaluable resource (https://sexchickengtex.farmgtex.org) that not only advances chicken genetics and genomics but also offers novel insights into the evolution of sexual dimorphism in vertebrates.

**Fig. 1.**
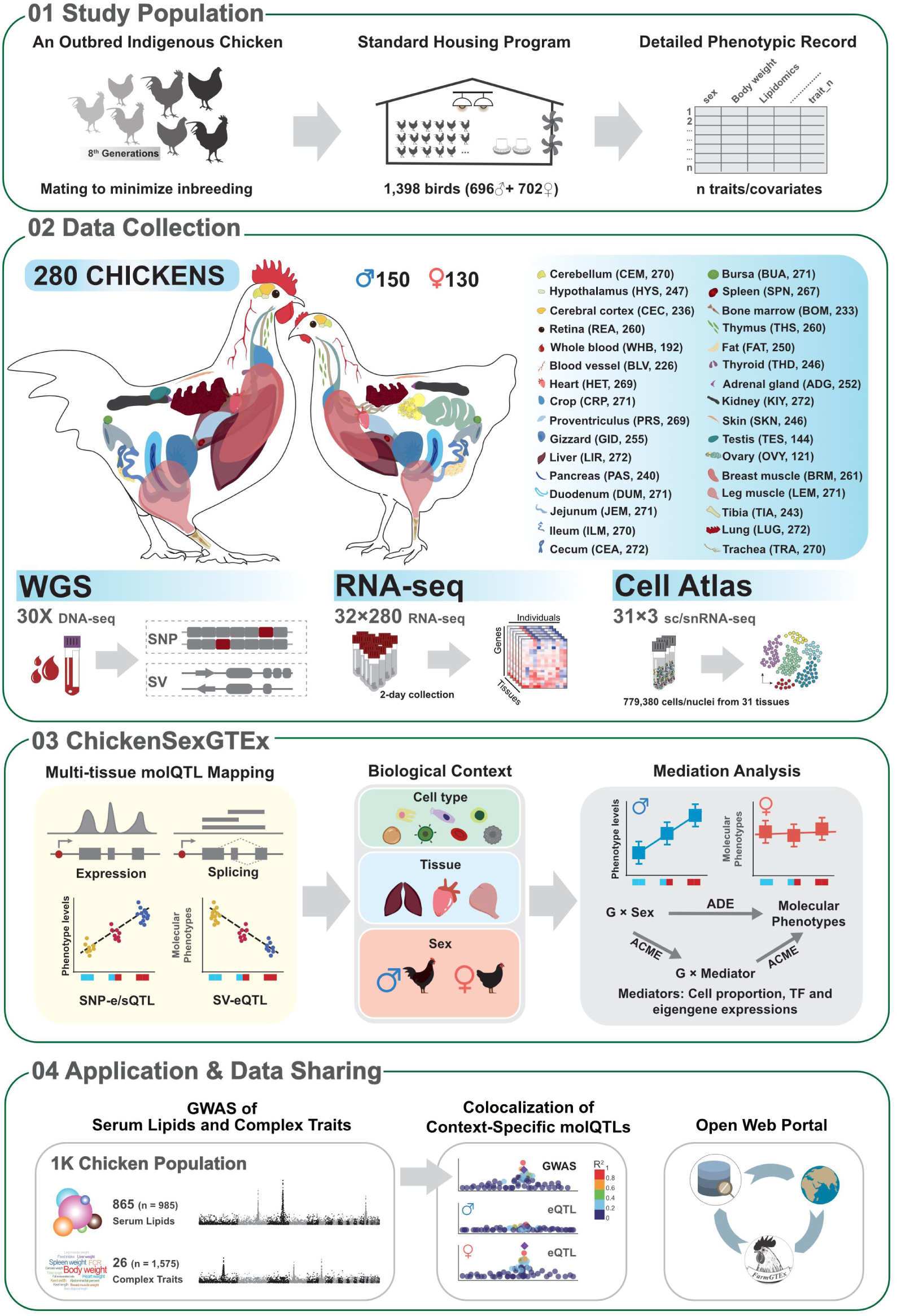
Experimental design and analytical workflow of the ChickenSexGTEx resource.

### Overview of the ChickenSexGTEx resource

In total, we analyzed the 280 high-depth WGS datasets, identifying 18,313,383 SNPs, 2,005,154 short insertion and deletion (Indels), and 137,596 SVs (length > 50bp). The SV catalog was dominated by insertions (INS, 110,882) and deletions (DEL, 26,207), with fewer inversions (INV, 131) and duplications (DUP, 376) (**Fig. 2A, fig. S1A**). For downstream *cis*-expression/splicing quantitative trait loci (e/sQTL) mapping, we focused on approximately 15 million common variants (minor allele frequency, MAF > 0.05), including 13,491,167 SNPs, 1,408,682 Indels, and 113,607 SVs (**Fig. 2A**). On average, all 25,358 annotated genes in the chicken genome were flanked by 27,350 SNPs, 2,717 Indels, and 254 SVs within their *cis*-regulatory windows (±1Mb of the transcript start site, TSS) (**fig. S1B**). Frequency and LD decay pattern of genetic variants were highly consistent between sexes (**Fig. 2B**), with LD falling below 0.1 within 1.5 kb for SNPs and 1.3 kb for SVs (**Fig. 2C and fig. S1C**). Notably, nearly half of the identified SVs (49.2%) showed weak linkage (LD < 0.2) with nearby SNPs or Indels (**Fig. 2D**), suggesting that the inclusion of SVs can capture genetic variation that would be missed by SNP-only analyses.

**Fig. 2.**
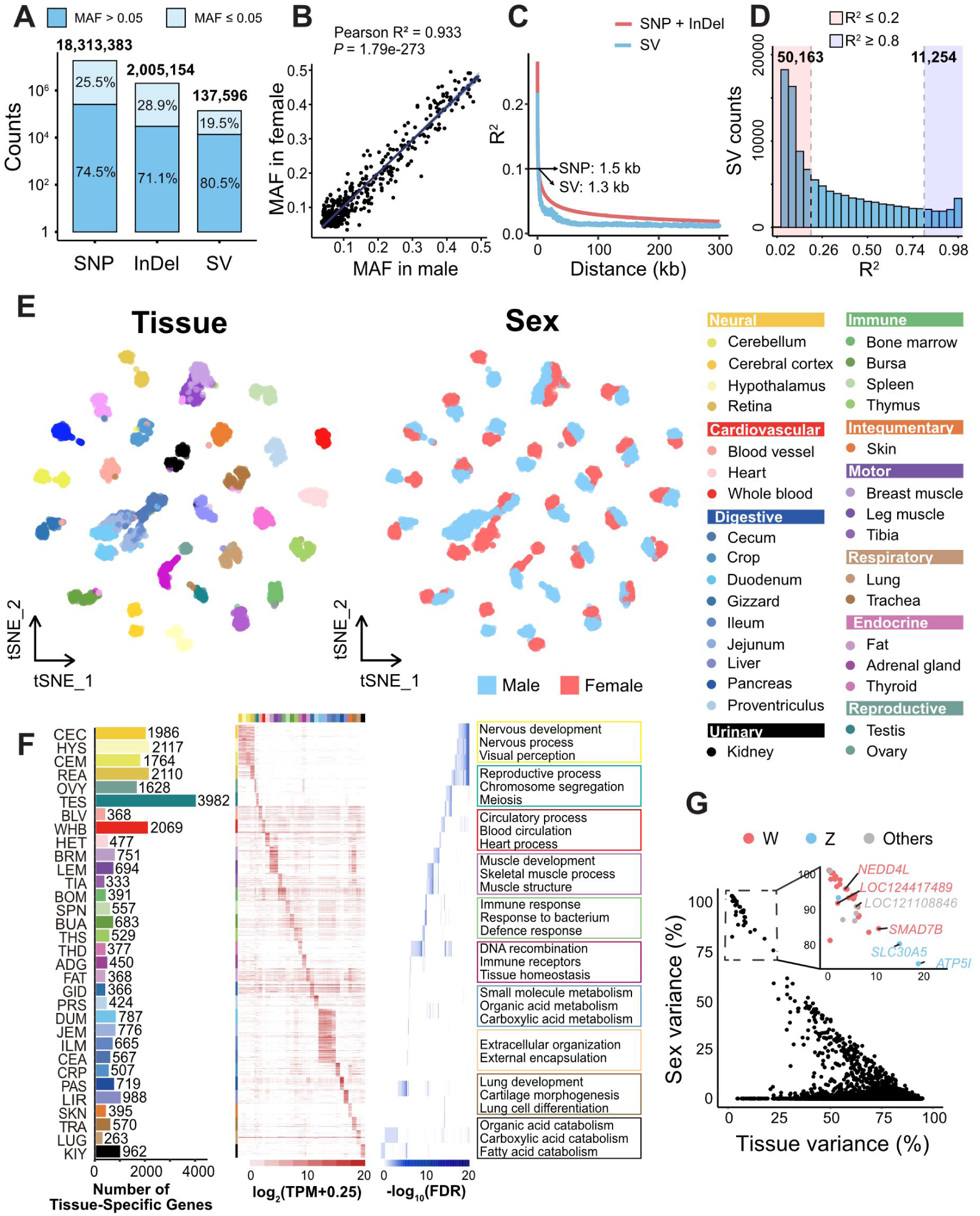
Genetic variant profiling and transcriptomic landscape across 32 tissues of ChickenSexGTEx cohort. **(A)** Overview of genetic variants identified via 30×whole-genome sequencing across 280 individuals. SNP: single nucleotide polymorphism; InDel: short insertion and deletion; SV: structural variants. **(B)** Comparison of minor allelic frequency (MAF) of common SNPs between male and female subpopulations, demonstrating the concordant genetic backgrounds. **(C)** Linkage disequilibrium (LD) decay patterns for SNP/InDel and SV, measured as pairwise correlation coefficient (R^2^) against genomic distance. **(D)** Counts of SVs that are strongly linked (R^2^ > 0.8) or largely unlinked (R^2^ < 0.2) to nearest SNPs across the genome. **(E)** t-distributed stochastic neighbor embedding (t-SNE) dimensionality reduction (*119*) of 7,969 RNA-seq samples, projected using the top 5,000 variably expressed genes, colored by tissue (left panel) and sex (right panel), respectively. **(F)** Identification and characterization of tissue-specific genes. Panels (from left to right) depict expression levels, statistical significance of tissue specificity, and enriched biological pathways, respectively. Tissue abbreviations are referred to Fig. 1. (**G)** Variance partitioning of gene expression by sex and tissue effects, quantified using ANOVA. Each point represents one gene, with axes showing the proportion of variance explained by each factor. W, Z and Others denote W chromosome, Z chromosome and autosomes, respectively.

For molecular phenotyping, we processed 8,265 RNA-seq samples across 32 primary tissues. After stringent quality control, we retained 7,969 high-quality transcriptomes from 280 individuals, with 274 chickens represented by more than 25 tissues each (**fig. S2A, data S1**). We achieved an average depth of 24.74 million clean reads per sample with a 92.16% mapping rate, enabling the robust detection of 22,473 expressed genes (transcript per million, TPM > 0.1) and 15,028 alternative splicing events (**fig. S2B**). Gene expression profiles were primarily segregated by tissue identity, with sex serving as a secondary driver of variation. In contrast, alternative splicing patterns clustered strictly by tissue, exhibiting no pronounced sexual dimorphism (**Fig. 2E, fig. S2C and D**). On average, we identified 925 tissue-elevated expressed genes per tissue, which recapitulated specialized physiological functions, ranging from neural morphogenesis in brain to sperm meiosis in the testis (**Fig. 2F**).

Analysis of variance (ANOVA) confirmed that tissue and sex explained significantly higher proportions of variance in gene expression (80% and 0.8%, respectively) compared to splicing (38% and 0.1%), whereas splicing exhibited stronger individual-specific effects (4.5%) than gene expression (3.7%) (fig. S3A). Genes whose expression were predominantly driven by sex (sex explained variance > 50%) were significantly enriched on sex chromosomes and involved in insulin metabolism and ion transport (**Fig. 2G and fig. S3B**). This is to be expected, with the incomplete dosage compensation in birds resulting in males having an average of 30% increased gene expression for genes on the Z chromosome (excluding those in the Pseudo Autosomal Regions)(*15*). Conversely, genes with high individual-specific splicing variation were associated with core cellular housekeeping functions, such as ribosomal and cytoskeletal components (**fig. S3C**). Furthermore, we evaluated the impact of 16 metadata—including sampling time, growth rates, body composition, and organ weights—on the chicken transcriptome (**data S2**). Notably, sampling time was a major driver of gene expression variance (mean 2.90%) and genes highly sensitive to sampling time included the core circadian clock genes such as *CIART*, *PER2*, *PER3*, and *ARNTL* (*26, 27*) (**data S3 and fig S3D**).

### Discovery of regulatory effects and tissue-sharing patterns

To investigate the genetic architecture governing the chicken transcriptome, we first estimated the *cis*-heritability (*cis-h^2^*) of gene expression and alternative splicing across all 32 tissues, yielding means of 0.29 and 0.08, respectively (**fig. S4**). While *cis-h^2^* was highly correlated between expression and splicing across tissues (Spearman’s ρ = 0.78, *P* = 1.15×10^−06^), we identified tissue-specific discordances in genetic control, particularly for neural tissues, such as retina, hypothalamus and cerebral cortex (**Fig. 3A**). We next performed *cis*-e/sQTL mapping, identifying an average of 11,599 genes possessing at least one eQTL (hereafter referred to as ‘eGenes’) and 5,235 genes possessing at least one sQTL (hereafter referred to as ‘sGenes’) per tissue (**Fig. 3B and fig. S5**). The discovery power was strongly driven by *cis-h^2^* across tissues (f**ig. S4C and fig. S4D**). Conditional analysis revealed 48% of eGenes and 28% of sGenes harbored at least two independent QTL (**Fig. 3C, table S5**), a finding corroborated by Bayesian fine-mapping analysis (*28*) (**fig. S6B and C**). Notably, fine-mapping yielded 175,807 high-confidence credible sets (*PIP* > 0.8) of eQTL, among them 59,475 (33.83%) were resolved to a single causal variant. Genes with more independent eQTL tended to have higher *cis-h^2^*, lower expression levels, and reduced sequence conservation. Furthermore, compared to primary eQTL (the SNP with the most significant level on any given eGene), non-primary ones tended to have lower MAF, smaller effect size, and farther distance from the TSS (**fig. S7**). As expected, e/sQTL were significantly enriched in TSS, transcriptional ending site (TES), and functional regulatory elements (**fig. S6A and D, fig. S8D, fig. S9A**). The robustness of our QTL was further validated by allele-specific expression (ASE) analysis, which showed high correlation with eQTL effect sizes (mean *r* = 0.91) (**fig. S10A and B**). Compared to the ChickenGTEx pilot phase,(*25*) our refined cohort provided a 14- to 18-fold increase in QTL discovery, significantly higher replication rates, and superior resolution for fine-mapping variants (**fig. S8, A to C and fig. S10, table S5**).

**Fig. 3.**
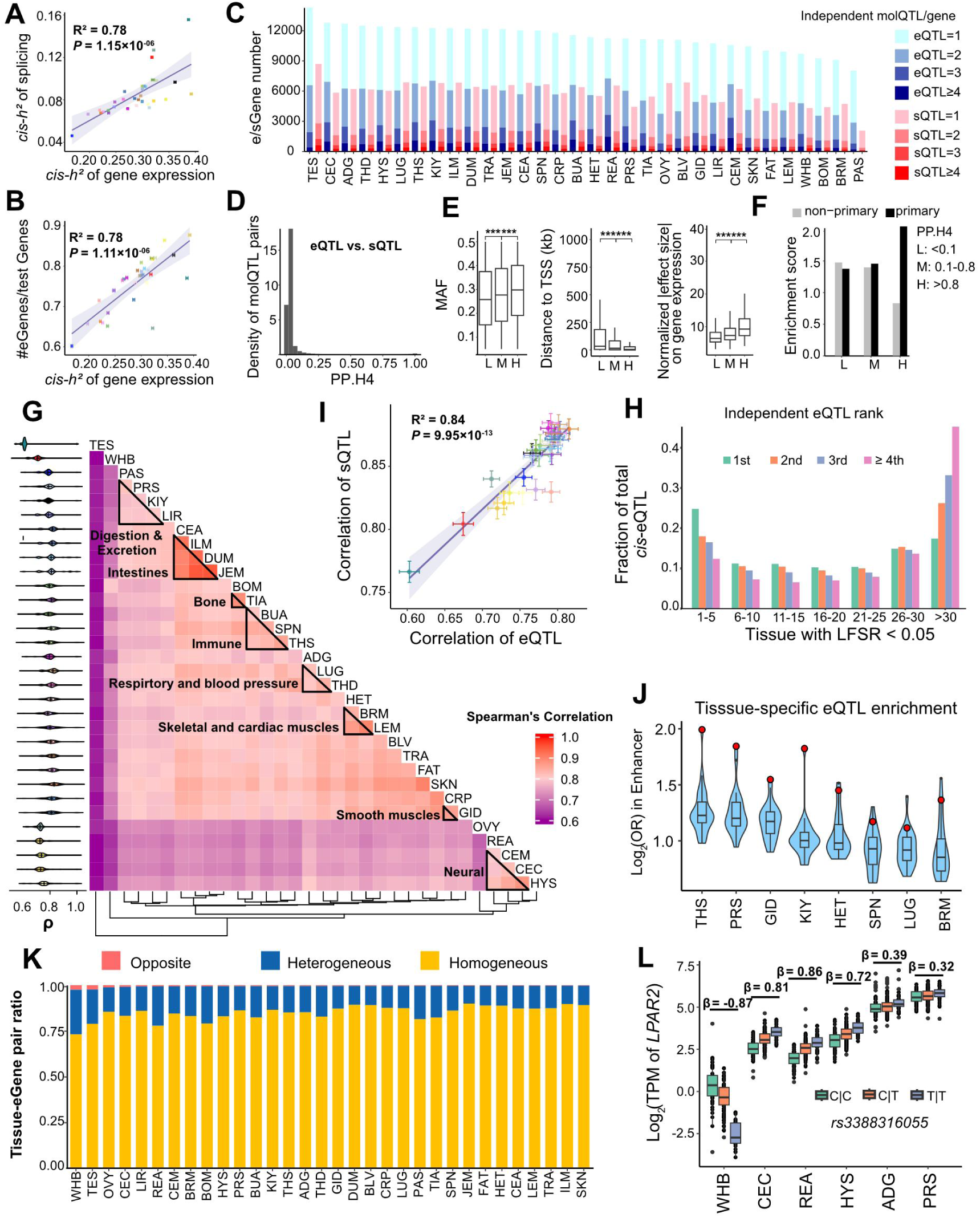
Multi-tissue molQTL mapping and tissue-sharing patterns. **(A)** *Cis*-heritability estimates for gene expression and alternative splicing across 32 distinct tissues. Point color denotes tissue type, consistent with the color key provided in Fig. 2E. **(B)** Correlation between the proportion of eGenes and *cis*-heritability of gene expression per tissue. Point color corresponds to tissue type, consistent with the color key provided in Fig. 2E. **(C)** Distribution of eGenes and sGenes categorized by the counts of conditionally independent eQTL or sQTL per gene (1, 2, 3, or ≥4). Tissue abbreviations are provided in Fig. 1. **(D)** Distribution of the posterior probability of colocalization (PP.H4) for eQTL and sQTL signals. **(E)** Minor allelic frequency (MAF), distance to transcription start site (TSS), and effect size of eQTL, stratified into low (L), median (M) and high (H) colocalization probability with sQTL. Asterisks indicate significance from ANOVA comparison. *** *P* < 0.001. **(F)** Enrichment of primary and non-primary eQTL signals across low (L), median (M) and high (H) colocalization categories with sQTL. **(G)** Pairwise tissue similarity of eQTL effect sizes, shown as Spearman’s correlations (left: Distribution of correlation values per tissue; bottom: hierarchical clustering of tissues by regulatory similarity). Tissue abbreviations are provided in Fig. 1. **(H)** Correlation between the effect size correlations of eQTL and sQTLs in each tissue. Points represent the mean effect size correlation of molQTL, and error bars denote the full range of correlation values. Colors indicate tissue type, as in Fig. 2E. **(I)** Fraction of eQTL shared across increasing numbers of tissues (LFSR < 0.05). The eQTL rank is determined by stepwise conditional analysis. **(J)** Enrichment of tissue-specific eQTL (detected in only one tissue) within annotated enhancers across different tissues. Red points represent the enrichment score for the matched tissue. **(K)** Classification of lead eQTL regulatory effects as opposite (red), heterogeneous (blue), or homogeneous (yellow) across tissues. **(L)** Tissue-specific regulation of variant *rs3388316055* on *LPAR2* expression, demonstrating opposite allelic effects in whole blood compared to neural and endocrine tissues.

We then investigated whether gene expression and splicing share a common genetic basis. Colocalization analysis revealed a striking divergence: 71.82% of eQTL showed no evidence of colocalization with sQTL (PP.H4 < 0.1 and PP.H3 > 0.8), with a negligible mean LD of 0.07 (**Fig. 3D and fig. S9B**). Only a small fraction of variants (1.69%, PP.H4 > 0.8) exhibited pleiotropic effects on both molecular phenotypes. These pleiotropic variants were characterized by higher MAFs, larger effect sizes, and a higher enrichment in promoters (**Fig. 3E**). Furthermore, 70.34% of pleiotropic variants were primary eQTL (**Fig. 3F, fig. S9C and D**). Collectively, these results suggest that for the majority of genes, the genetic control of total gene expression is decoupled from the regulation of alternative splicing.

Using mashR to model tissue-sharing dynamics (*29*), we identified a U-shaped distribution of regulatory effects, where variants were either highly ubiquitous or strictly tissue-specific (*30*) (**Fig. 3H**). Tissues with similar physiological ontologies such as immune or neural systems clustered together based on their regulatory effect sizes, a pattern consistent across both eQTL and sQTL (Spearman’s ρ = 0.84, *P* = 9.95 ×10^−13^) (**Fig. 3G**). While 10.78% of eQTL were shared across all 32 tissues, 18.01% were active in only a single tissue (**Fig. 3H**). These tissue-specific regulatory variants were markedly enriched at TSS and exhibited larger effect sizes and stronger enhancer enrichment in their corresponding tissues (**Fig. J and fig. S11A**). In contrast to primary variants, non-primary variants were more likely to be shared across tissues but showed lower enrichment in promoters (**fig. S11B**). Although sQTL followed similar tissue sharing patterns, they exhibited a higher overall degree of tissue-sharing (86.60% vs. 71.99% in eQTL) (**fig. S11C and D**). Notably, tissue-specific variants were more likely to be pleiotropic, characterized by high colocalization between eQTL and sQTL (PP.H4 > 0.8), a phenomenon predominantly enriched in reproductive tissues such as testis and ovary (**fig. S9, E to G**).

To investigate tissue-specific regulatory variants, we compared effect size of primary e/sQTL across tissues, classifying them as homogeneous (concordant direction with overlapping 95% confidence intervals, CIs), heterogeneous (concordant direction with non-overlapping 95% CIs), and opposite (different directions). Compared to eQTL (14.97%), sQTL showed a lower proportion (14.08%) of heterogeneous and opposite effects (**Fig. 3K and fig. S11E**). Consistent with mashR results above, whole blood (2.52%), testis (2.41%) and ovary (1.23%) had a relatively high fraction of directionally opposite effects compared to other tissues, whereas skin showed the highest directional stability, with only 0.32% of eQTL being opposite (**Fig. 3K**). While rare, these directionally opposite variants might provide novel insights into complex physiological trade-offs. For example, the T allele of *rs3388316055* is associated with decreased expression of *LPAR2* in whole blood but significantly increased expression in the cerebral cortex, retina, and adrenal gland (**Fig. 3L and fig. S12**). Given the dual roles of *LPAR2* in modulating vascular integrity and oncogenesis, these antagonistic effects likely reflect highly specialized tissue-specific regulatory environments. Such regulatory pleiotropy may explain how a single locus exerts divergent or even conflicting influences on complex traits across different physiological systems (*31, 32*).

### Single-cell deconvolution reveals cell-type-dependent regulatory landscapes

While bulk transcriptomics provides an overview of gene regulation, expression profiles are a function of the underlying cellular landscape. To bridge this resolution gap, we established a comprehensive chicken single-cell atlas, comprising 779,380 cells/nuclei across 31 tissues, meticulously matched to our bulk RNA-seq cohort. By leveraging this atlas for computational deconvolution, we conducted cell-type interaction eQTL (cieQTL) mapping (**Fig. 4A, fig. S13A and B**). Based on canonical marker genes (**data S4**), we identified 57 broad cell types across 8 major lineages: immune, endothelial, epithelial, muscular, reproductive, hematopoietic, neural, and mesenchymal (**Fig. 4B, fig. S13C, fig. S14**). To achieve a higher degree of granularity, we manually annotated 127 distinct cell types (mean 12.77 per tissue), providing a high-definition cellular map of the chicken (**fig. S15 to 22**).

**Fig. 4.**
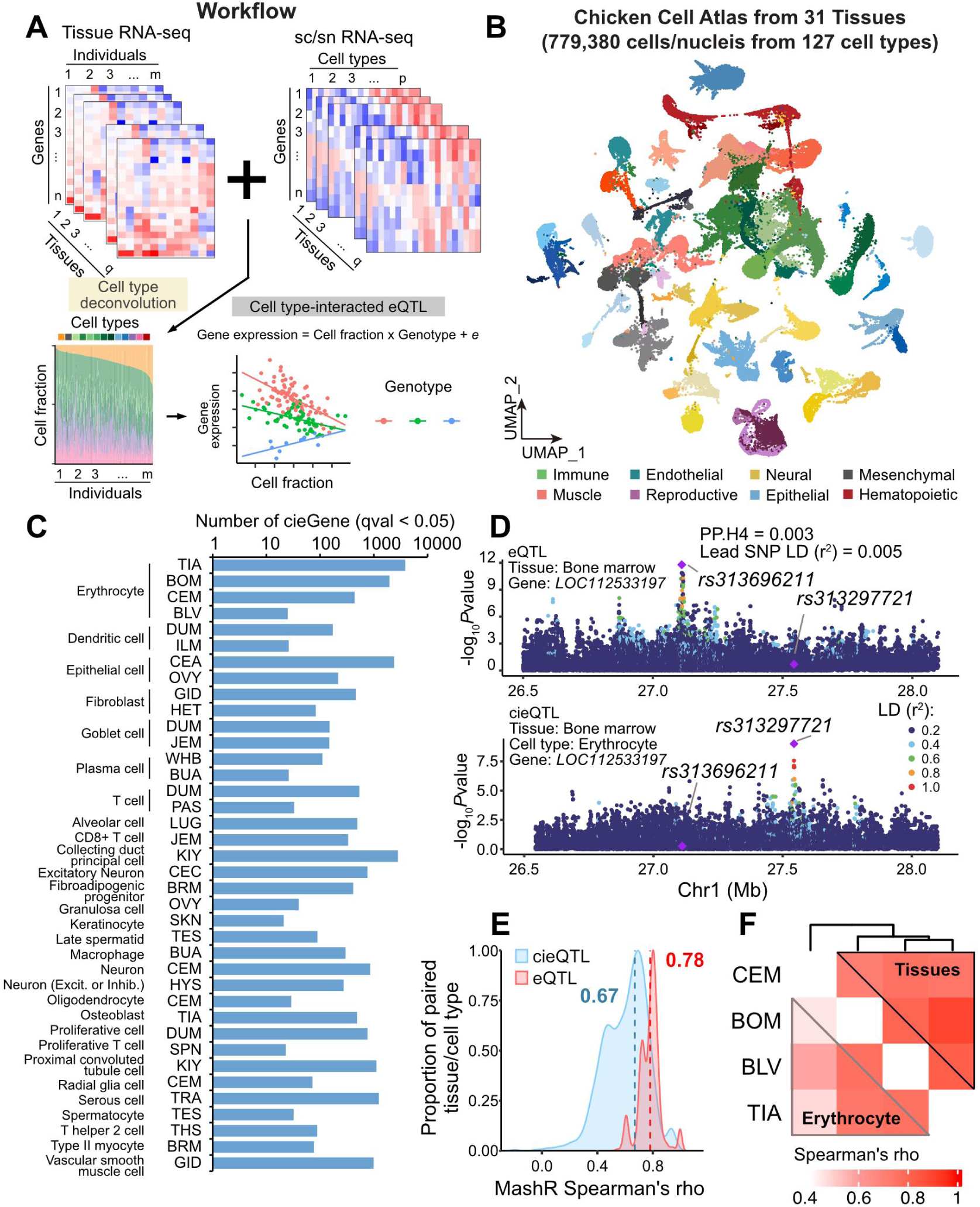
Single-cell atlas and cell-type-interaction regulatory effects. **(A)** Analytical workflow integrating single-cell/single-nucleus RNA-seq with bulk RNA-seq for estimating cell fraction deconvolution and mapping cell-type-interaction eQTL (cieQTL) mapping. **(B)** Uniform manifold approximation and projection (UMAP) visualization of 779,380 single cells/nuclei across 31 tissues, clustered into 8 distinct cell lineages. **(C)** Number of cieGenes identified across 38 distinct tissue-cell type pairs. Tissue abbreviations are provided in Fig. 1. **(D)** An example of discordant genetic regulation between a cieQTL and a steady bulk steady-state eQTL in bone marrow: the erythrocyte cieQTL (*rs313297721*) for *LOC112533197* shows no colocalization (PP.H4 = 0.003) with bulk bone marrow bulk eQTL (*rs313696211*). **(E)** Sharing patterns of cieQTL and bulk eQTL across matched pairs of cell types and tissues. **(F)** Spearman correlation of genetic effect sizes for for erythrocyte-specific cieQTL (lower left panel) and bulk eQTL (upper right panel) across cerebellum, bone marrow, blood vessel, and tibia tissues. Tissue abbreviations are provided in Fig. 1.

We utilized this cell atlas to deconvolute the cell fractions of all 7,969 bulk RNA-seq samples, revealing a dynamic and heterogeneous landscape of cellular compositions (**fig. S23 to 30, data S5**). While 76 out of 127 cell types were strictly tissue-specific, others—such as endothelial and blood cells—were ubiquitous but fluctuated significantly in proportion (**fig. S31A and B**). By integrating 28 collected metadata, we observed that 22 were significantly associated with the shifts of intra-tissue cellular proportions across individuals (**fig. S32A**). Notably, muscle accretion was primarily associated with cellular proportions in the gastrointestinal tract, such as goblet cells in the jejunum and proliferative cells in the duodenum—cell types essential for nutrient digestion and absorption. For subsequent analysis, we kept 79 tissue-cell pairs with median fractions exceeding 10% (**fig. S31C**). After performing GWAS of cell type composition, we detected 58 out of 79 tissue-cell pairs that were significantly associated with at least one genomic variant (cell-fraction QTL, hereafter referred as to cfQTL) (**Fig. 4C, fig. S32B and C**). In total, we detected 119 cfQTL (*P* < 5 ×10^−8^), among which 84 (70.59%) were colocalized with eQTL (PP.H4 > 0.8). Most of cfQTL were colocalized with eQTL in blood vessel (21.01%) and kidney (10.92%) (**fig. S33**). Particularly, 96.43% of cfQTL were colocalized with eQTL from a different tissue, indicative of a widespread cross-tissue regulatory network governing cell proportion. For instance, the variant *rs1057666964*, which was associated with Purkinje cell proportion in the cerebellum, exclusively colocalized with an eQTL of *MYO1D* in the lung, suggesting that the genes influencing the lung contrasting functions may be also connected with the cerebellum neuronal integrity (*33, 34*). In contrast, *rs732753151* and *rs316114259*—associated with dendritic cell proportions in the ileum and duodenum, respectively—colocalized with eQTL for *BLB2,* a critical avian immune gene (*35, 36*), across 13 and 26 tissues (PP.H4 = 0.99) (**fig. S34A**).

After performing cieQTL mapping, we identified 10,937 cieGenes containing at least one cieQTL across 42 cell types in 29 tissues (**Fig. 4C and fig. S34B**). Notably, 53.61% of these cieQTL showed no evidence of colocalization with bulk eQTL (PP.H3 > 0.8) (**fig. S34C**), indicating that a portion of the regulatory genome is only active within specific cellular contexts. This is exemplified by *LOC112533197* in the bone marrow, where the regulatory signal is strictly dependent on erythrocyte abundance and remains masked in bulk tissue analyses (**Fig. 4D**). Furthermore, cieQTL exhibited significantly higher tissue-specificity than standard eQTL (**Fig. 4E, F**). Overall, 37.16% of cieQTL were active in only a single tissue-cell pair, compared to just 18% of standard eQTL (**fig. S34D**). A compelling case is the cieQTL *rs317111159*, which regulates the expression of *TRPC6* gene, a gene linked to Ca2+ leakage in human erythrocytes (*37, 38*). While this variant initially appeared to be a broad, tissue-shared eQTL across the blood vessel, cerebellum, and tibia, its regulatory activity was exclusively confined to erythrocytes within the vascular environment (**fig. S35**). These results demonstrate that resolving cellular heterogeneity can uncover latent, cell-type-specific genetic architectures that are otherwise obscured in traditional tissue-level investigations.

### Multi-layered architecture of sexual dimorphism in chicken

We systematically assessed differences in gene expression, co-expression networks, and cellular composition between sexes across 30 tissues (excluding gonads) (**Fig. 5A**). Our analysis revealed a pervasive sex-biased gene expression, identifying 16,854 (65.56% of all 25,708 annotated genes) genes with differential expression levels between sexes, referred to as sex-biased genes (sbGenes), in at least one tissue. The magnitude of sexual dimorphism was most pronounced in the tibia, bursa, and hypothalamus, while intestinal tissues exhibited the highest transcriptomic similarity between sexes (**Fig. 5B and data S6**). The majority (94.59%) of sbGenes were on autosomes, and the number of genes with upregulated expression in male (male-biased genes, mbGenes) were similar to those of female (female-biased genes, fbGenes) (**Fig. 5B**). In contrast to the chromosome-wide X-inactivation in mammals, we observed that 902 Z-linked genes remained male-biased across nearly all tissues, reflecting incomplete dosage compensation(*13*) (**Fig. 5B, fig. S36A**). Intriguingly, we identified a specific region on Z chromosome (ChrZ: 27-28Mb) that was significantly enriched with fbGenes across all 30 tissues (**fig. S36B**). This region overlaps with the male hypermethylation (MHM) domain, providing robust evidence for a localized, epigenetically regulated dosage compensation mechanism in birds (*14, 15*). Functionally, the sex-biased transcriptome exhibited a clear division of labor: mbGenes were significantly enriched in musculoskeletal development, whereas fbGenes dominated lipid metabolism and immune pathways (**fig. S36C**). In the tibia, for instance, the male-biased expression of *IHH*—a master regulator of chondrocyte maturation—supports the more robust skeletal growth observed in males. Conversely, the female-biased expression of *LYG2* underscores a superior antimicrobial defense profile in females (*39, 40*) (**Fig. 5C**). Collectively, these findings provide a molecular roadmap explaining why male birds prioritize skeletal robustness, whereas females invest more heavily in immunological responsiveness and metabolic efficiency (*41*). Beyond individual genes, we examined higher-order transcriptomic organization through co-expression network analysis. In total, we detected 672 co-expression modules across all tissues, of which 405 (60.2%) were significantly associated with sex (**Fig. 5B and data S7**). For instance, in the hypothalamus, we identified a female-biased module enriched for neuro-immune homeostasis and a male-biased module linked to circadian rhythms and growth axes (**Fig. 5D and fig. S37A**). These results suggest that sexual dimorphism in the avian “master regulator” (the hypothalamus) is achieved through distinct physiological strategies: females prioritize immune-neuronal stability (*42*), while males integrate growth and stress responses (*43, 44*).

**Fig. 5.**
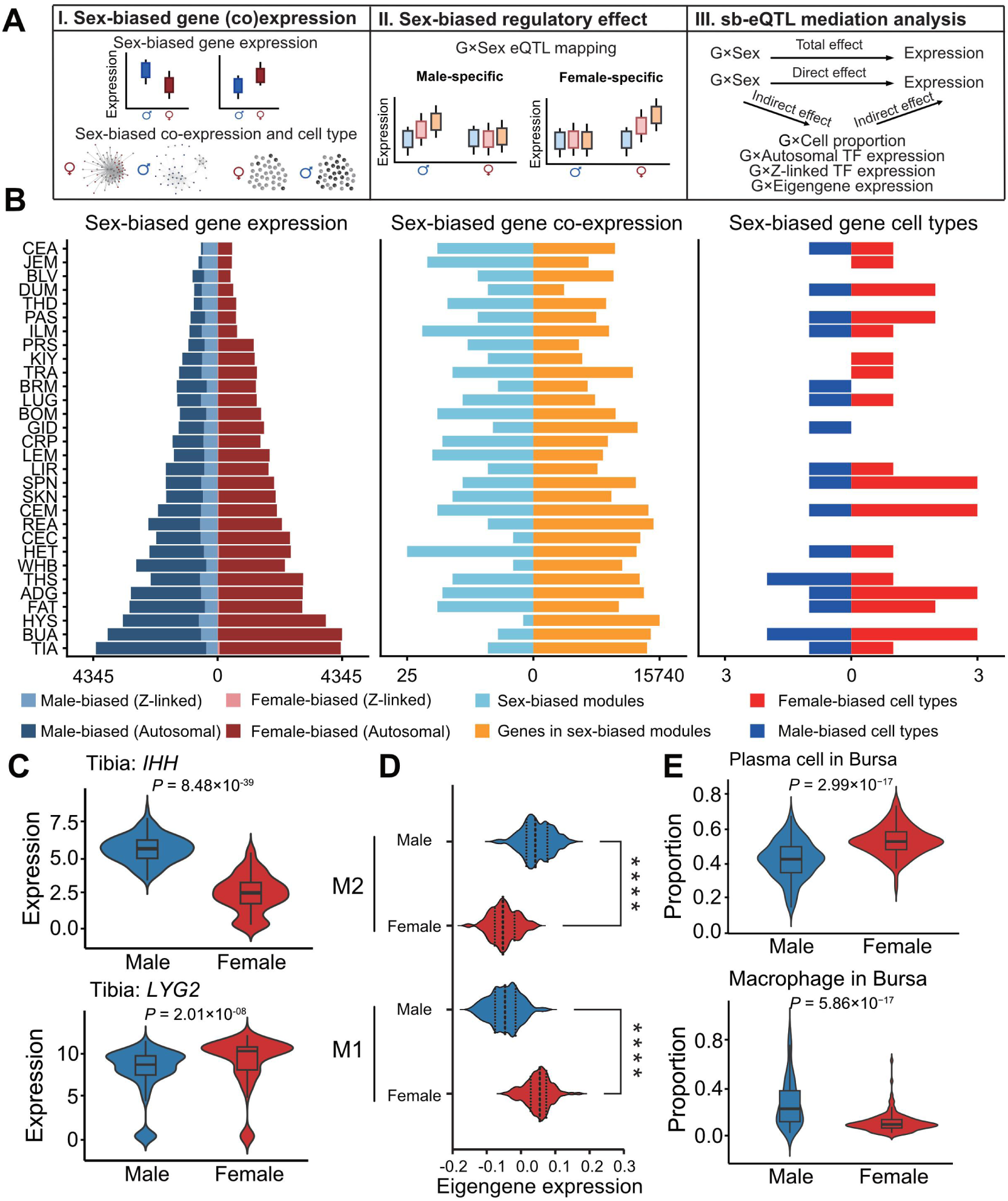
Differences in gene expression, co-expression networks, and cell proportions between sexes across 30 chicken tissues. **(A)** Schematic analytical workflow comprising three core modules: (I) characterization of sex-biased gene expression, co-expression networks, and cell type proportions; (II) identification of sex-biased genetic regulatory effects (sb-eQTL); (III) mediation analysis of sb-eQTL. **(B)** Summary of sex-biased molecular and cellular features across 30 tissues, including sex-biased gene expression (sb-Genes) (left), sex-biased co-expression networks (middle), and sex-biased cell types exhibiting sex-dimorphic abundance (right). Tissue abbreviations are provided in Fig. 1. **(C)** Expression profiles of male-biased gene (*IHH*) and female-biased gene (*LYG2*) in tibia tissue between males and females. **(D)** Eigengene expression levels of sex-biased co-expression modules, M1 and M2, in the hypothalamus, stratified by sexes. **(E)** Comparison of cell proportion in the bursa between male and female chickens, illustrating a male-biased increase in macrophages and a female-biased increase in plasma cells

We identified 30 cell types exhibiting significant sex differences in abundance across 19 tissues, referred to as sex-biased cell types (**Fig. 5B, and data S8**). Notably, plasma cells were consistently overrepresented in females across lymphoid tissues (**fig. S37C**), whereas macrophages were male-biased in the bursa, suggesting a fundamental skew toward humoral immunity in females and innate surveillance in males (*45, 46*) (**Fig. 5E**). Crucially, correcting for these cell-type proportions significantly reduced the number of detected sbGenes, demonstrating that a substantial fraction of observed “sex-biased expression” is actually a proxy for underlying shifts in cellular architecture between sexes (**fig. S37B**) (*22, 47*).

### Genetic basis of sex dimorphic regulation

To dissect the genetic component of sex dimorphism, we first compared *cis-h^2^* of gene expression between male and female. While *cis-h^2^* was broadly correlated across tissues (average Pearson’s *r* = 0.74), the lowest correlations were observed in fat and blood vessel (**fig. S38**). We then applied a Genotype×Sex (G×Sex) interaction model to identify sex-biased regulatory effects across tissues. Despite the presence of 16,854 differentially expressed genes, only 340 genes (sb-eGenes) were associated with sex-biased regulatory variants (sb-eQTL) (**Fig. 6A and fig. S39A**). Notably, we found a negligible overlap and weak correlation between sbGenes and sb-eGenes across tissues (**Fig. 6A and fig. S39B**). We further estimated effect sizes of sb-eQTL in males and females separately, revealing 194 sex-specific sb-eQTL (only detected as sb-eQTL in one sex) (**Fig. 6B**). For example, in the tibia, the myosin-encoding gene *MYO5C* expression was only significantly associated with *rs731939689* in male (**Fig. 6C**). Meanwhile, 161 sb-eQTL exhibited concordant allelic directions but different effect sizes between sexes (**Fig. 6D and fig S39C**). For instance, *GART*, a gene widely implicated in the biosynthesis of muscle flavor compounds, inosine monophosphate (*48*), demonstrated a significantly larger effect size in female than in male adipose (**Fig. 6E**). We further found 94 sb-eQTL with reverse allelic effects between sexes (**Fig. 6D, fig. S39C, data S9**), which displayed significantly smaller effect sizes than sb-eQTL with concordant effect (**fig. S39D and E**). Such genes are particularly interesting from a perspective of the resolution of sexual conflict (*49–53*). In this case, sexual conflict refers to diverging selection for specific phenotypes between males and females (for example if it is beneficial for males to have higher expression of a particular gene and beneficial for females to have lower expression of a particular gene). Regulatory evolution to modify the response in different sexes appears to be a highly efficient method to resolve such sexual conflicts (*54*). Out of 340 sb-eGenes, 306 were found in single tissues, and only 6 shared in more than five tissues (**fig. S39F**). For instance, *ZBTB11*, a transcription factor having housekeeping roles in maintaining the mitochondrial function in humans (*55*), exhibited male-specific genetic regulation across 23 tissues (**fig. S40**). This suggests a male-specific genetic “program” for energy metabolism to support higher muscle accretion and physical activity (*56–58*).

**Fig. 6.**
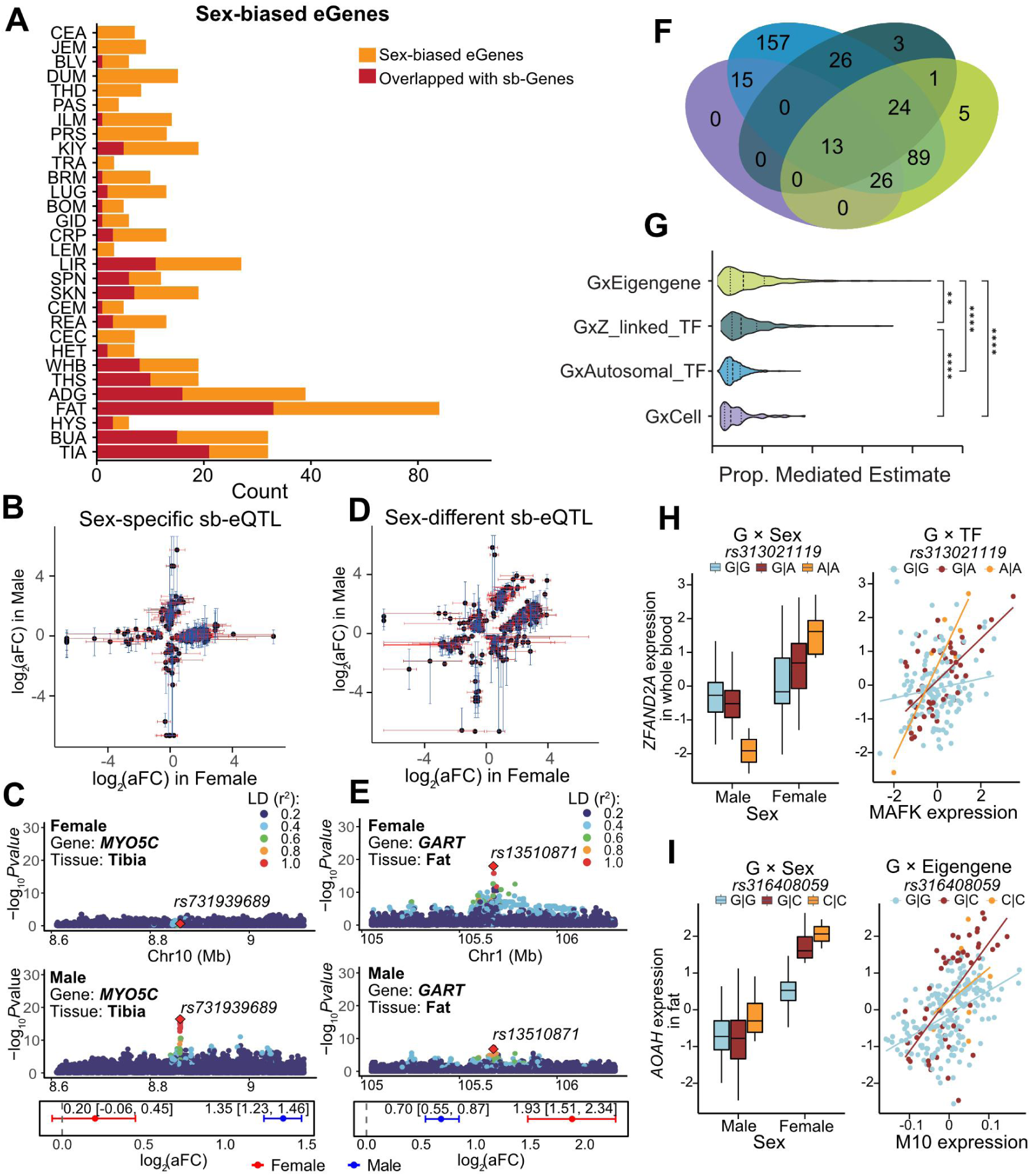
Sex-biased genetic regulation and mechanistic mediation analysis. **(A)** Summary of sb-eGenes identified across 30 tissues, highlighting the overlap between sb-eGenes and sb-Genes. Tissue abbreviations are provided in Fig. 1. **(B)** Effect size estimates for sex-specific eQTL derived from sex-stratified analysis. Points denote the mean effect size (log_2_FC), and error bars represent the 95% confidence intervals. Sex-specific eQTL were defined as loci where the confidence interval of effect size overlaps with zero in one sex but not in the other. **(C)** Example of a male-specific sb-eQTL (*rs731939689*) that regulates *MYO5C* expression exclusively in the tibia. Bottom panel displays effect size estimates from male- and female-stratified populations, respectively. **(D)** Effect size estimates for sex-different and sex-reversed sb-eQTL derived from sex-stratified analysis. Sex-different sb-eQTL are loci with directionally concordant effects yet different magnitudes between sexes (Quadrants I and III), while sex-reversed eQTL are loci with directionally discordant effects between sexes (Quadrants II and IV). **(E)** Example of a sex-different sb-eQTL (*rs13510871*) that exerts a stronger regulatory effect on *GART* expression in female fat tissue than male fat tissue. **(F)** Venn diagram illustrating the mediation landscape of sb-eQTL, with contributions from co-expression modules, transcription factors (Z-linked or autosomal), and cell type proportions, where the colors of four contributions are consistent with the color key provided in Fig. 6G. **(G)** Distribution of proportion-mediated estimates across four mediator sets. Asterisks indicate significance from ANOVA comparison (\**P* < 0.05, \*\**P* < 0.01, \*\*\**P* < 0.001, and \*\*\*\**P* < 0.0001). **(H-I)** Examples of sb-eQTL mediation analsysis: (**H**) The *ZFAND2A* sb-eQTL mediated by the transcription factor *MAFK* in blood tissue; (**I**) The *AOAH* sb-eQTL mediated by the co-expression module M10 in fat tissue.

Finally, we identified the mediators of these G × Sex interactions by examining 30 sex-biased cell types, 699 sex-biased transcriptional factor (TF) genes, and 405 sex-biased co-expression modules (**data S10**). A total of 359 sb-eQTL (80%) were significantly mediated by at least one of three factors, with co-expression modules showing the strongest number of mediations, followed by Z-linked TFs (**Fig. 6F and G, fig. S41**). For instance, in whole blood, the sex-biased regulation of *ZFAND2A* was statistically mediated by *MAFK* expression, which is a key regulator of multiple hematopoietic lineages (*59*) (**Fig. 6H**). Likewise, in adipose tissue, the sb-eQTL of *AOAH*, an LPS-detoxifying lipase coding gene, was mediated by a co-expression module significantly enriched in immune-defense pathways (**Fig. 6I and fig. S41A**). Together, these results provide a holistic model where sexual dimorphism emerges not from isolated SNPs, but from a coordinated interplay between sex-linked genetic triggers, transcription factor networks, and dimorphic cellular landscapes.

### Structural variants associated with tissue-specific regulation and sexual dimorphism

While SNPs are the primary focus of most regulatory studies, structural variants (SVs) often exert more profound functional consequences due to their ability to disrupt or reorganize large genomic segments (*60, 61*). To quantify their contribution to the chicken regulatory landscape, we integrated 137,596 high-quality SVs into our *cis-h^2^* estimation models. The inclusion of SVs significantly increased the captured genetic variance in gene expression across all 32 tissues (Student’s t-test, *P* < 2.2 ×10^−16^) (**Fig. 7A**). Notably, 82.7% of genes showed higher *cis-h^2^* when SVs were accounted for, with the most dramatic gains occurring in genes with low SNP-based heritability (**fig. S42A**). While incremental improvements (1%–5%) were typical for ubiquitously expressed genes, substantial heritability shifts (>10%) were predominantly concentrated in tissue-specific genes (**Fig. 7B and fig. S42B**). These findings suggest that SVs are not merely auxiliary markers but are key determinants of tissue-specialized transcriptomic niches.

**Fig. 7.**
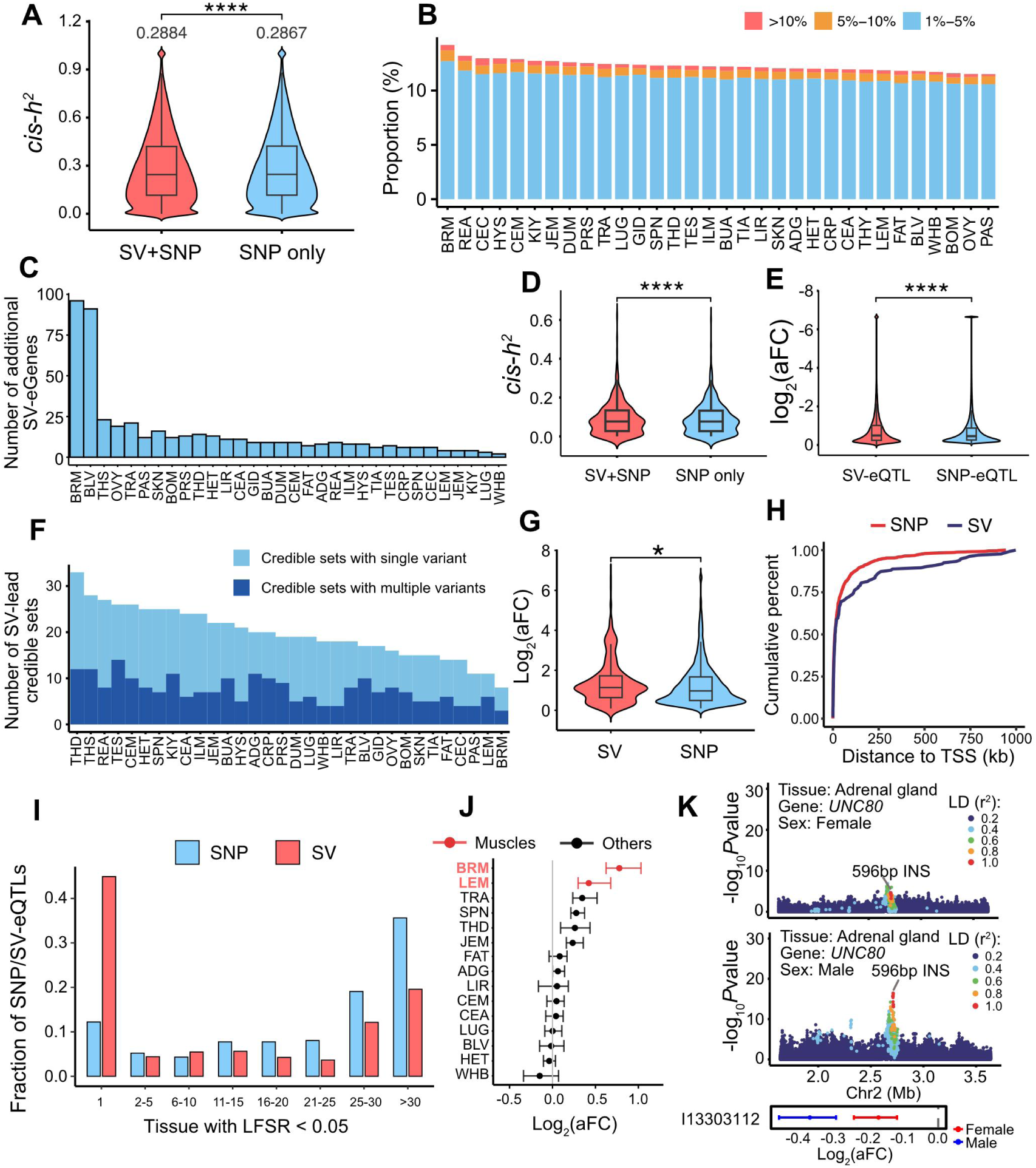
Regulatory impact of structural variants (SVs) on gene expression. **(A)** Comparison of gene expression *cis*-heritability (*cis*-*h*²) estimates from SNP-only models versus SNP+SV integrated models. **** represents the significance of *P* value < 0.0001. **(B)** Distribution of *cis-h²* gains attributable to SV inclusion in the model, stratified by tissue type. Tissue abbreviations are provided in Fig. 1. **(C)** Number of additional SV-eGenes identified upon integrating SVs across 32 tissues. **(D)** *cis-h²* estimates for the newly identified SV-eGenes, under SNP-only and SNP+SV integrated models. **** represents the significance of *P* value < 0.0001. **(E)** Comparison of down-regulatory effect sizes between SV-eQTL and SNP-eQTL. **** represents the significance of *P* value < 0.0001. **(F)** Number of 95% credible sets where an SV is the sole top causal variant or has a higher posterior probability than linked SNPs. **(G)** Effect size comparison between fine-mapped SV- and SNP-leading variants after matching for minor allele frequency. * represents the significance of *P* value < 0.05. **(H)** Cumulative distribution of genomic distances from fine-mapped SV- and SNP-leading regulatory variants to the TSS of their target genes. **(I)** Tissue-sharing patterns of SV-eQTL versus SNP-eQTL across 32 tissues. **(J)** Muscle-biased regulatory effect of a 66 bp insertion (INS) variant on *FSBP* gene expression. **(K)** Male-specific regulation of *UNC80* by a 596 bp INS in the adrenal gland.

We identified 15,013 genes associated with at least one SV (SV-eGenes), representing 73.1% of the total tested genes (**fig. S42C**). Although many SV signals were in high LD with neighboring SNPs, we uncovered 456 “SV-only” eGenes whose expression was exclusively regulated by SVs. These loci were particularly prevalent in metabolic tissues such as breast muscle and blood vessels (**Fig. 7C and fig. S42D**). These SV-only eGenes contributed significant additional heritability compared to SNP alone (**Fig. 7D**). For example, a 551 bp deletion was independently associated with the expression of *FGF9* in adipose tissue. Given that *FGF9* is a critical adipokine for energy metabolism and *UCP1* induction (*62*), this SV might represent a regulatory candidate for metabolic efficiency (**fig. S42E**). Notably, SVs exerted significantly more potent down-regulatory effect (Student’s t-test, *P* < 2.2 ×10^−16^) on gene expression compared to SNPs (**Fig. 7E**), likely due to their greater capacity to physically disrupt promoter-enhancer interactions or open reading frames. To detect causal effects of SVs from potential LD confounding effects on neighboring SNPs, we performed Bayesian fine-mapping (*28*) to identify 95% credible sets across all tissues (**Fig. 7F, data S11**). Although SVs were prioritized as the lead causal variants in a relatively small fraction of credible sets (1.48‰–4.32‰), they exhibited significantly larger effect sizes than MAF-matched SNP-eQTL (Student’s t-test, *P* = 0.025) (**Fig. 7G and fig. S42F**). These high-impact SVs were more frequently located in distal regions and exons rather than TSS, suggesting they may alter long-range enhancers (**Fig. 7H, fig. S42G and H**). A compelling example is a 388 bp deletion in the 5’UTR of *HHATL*, which exerts a dominant regulatory effect over SNPs in the kidney (**fig. S42I**).

A defining feature of SVs in our cohort was their heightened tissue-specificity relative to SNPs (**fig. S43A**). Specifically, 44.9% of SV-eQTL were strictly tissue-specific, with only 19.6% shared across more than 30 tissues. In contrast, SNP-eQTL showed an opposite pattern, being predominantly widely shared (35.6%) rather than tissue-specific (12.2%) (**Fig. 7I**). This suggests that SV-eQTL preferentially operate through tissue-specific mechanisms (**fig. S43B**). A tissue-specific example is a 66 bp INS that showed particularly strong effects (average *β* = 0.61) on *FSBP* expression exclusively in breast and leg muscle tissues (**Fig. 7J**), which was reported previously to be highly expressed in fast-growing broiler chickens (*63*). Furthermore, we identified 104 sex-biased SV-eQTL (sbSV-eQTL). The majority of them (96.2%) were tissue-specific, with the most pronounced enrichment in endocrine (adrenal gland) and immune (bursa) organs (**fig. S43C**). This again revealed the importance of sex-differential regulation in governing endocrine and immunological homeostasis. Remarkably, 87.50% (91) of these sex-dimorphic signals were unique to SVs and missed by SNP-only mapping, and 66.35% (69) of these sbSV-eQTL showed sex-specific regulatory effects. These variants primarily drove sex-biased effects on 23 lncRNA and 81 protein-coding genes (**fig. S43D**). For instance, a 596 bp insertion was associated with *UNC80* expression with a larger effect size in the male adrenal gland. Conversely, a 77 bp deletion modulated the phosphoprotein scavenger *MAGI2* more significantly in the female bursa (*64*) (**Fig. 7K and fig. S43E**). These findings demonstrate that SVs represent a distinct layer of the regulatory genome, playing a role in shaping the molecular boundaries between tissues and sexes.

### Sex-specific molecular architecture of the chicken lipidome

To investigate how the multi-tissue regulatory landscape translates into systemic physiological variation, we established a serum lipidomic metabolites GWAS (mGWAS) cohort of 985 chickens (525 male and 460 female) (**data S12 and S16**). By performing Bayesian colocalization analysis between molQTL (i.e., eQTL, sQTL, and cieQTL) and 279 independent GWAS loci associated with 865 serum lipids, we sought to elucidate the genetic cascade driving transcriptomic regulation toward metabolic output across five major lipid classes: sterols, fatty acyls, sphingolipids, glycerides, and glycerophospholipids. In total, 45.5% (127) of the mGWAS loci showed significant colocalization (PP.H4 > 0.8) with at least one molQTL (**Fig. 8A**). We observed distinct regulatory patterns across lipid classes: fatty acyls exhibited the highest proportion of colocalized loci (77.3%), followed by sphingolipids (49%) and glycerides (45%) (**Fig. 8A**). For example, a genome-wide significant locus on chromosome 11 (*rs731594651*) associated with serum acyl-carnitine (C2:0) levels colocalized with an eQTL of *VPS9D*1 in the cerebellum (PP.H4 = 0.91) (**Fig. 8B**). Given that V*PS9D1* activates Rab5-mediated endocytosis (*65*), this suggests a potential link between neuronal vesicular trafficking and systemic fatty acid metabolism (*66*). Furthermore, a GWAS signal of serum ceramide (Cer d18:1/18:1), a metabolite critical for mitochondrial redox balance (*67*), colocalized with a sQTL of *CD36* in the kidney (PP.H4 = 0.91) (**Fig. 8C**). Since *CD36* is a master regulator of long-chain fatty acid uptake and ceramide recruitment (*68, 69*), this finding prioritizes *rs312755963* as a key variant coordinating systemic lipid homeostasis through tissue-specific isoform regulation.

**Fig. 8.**
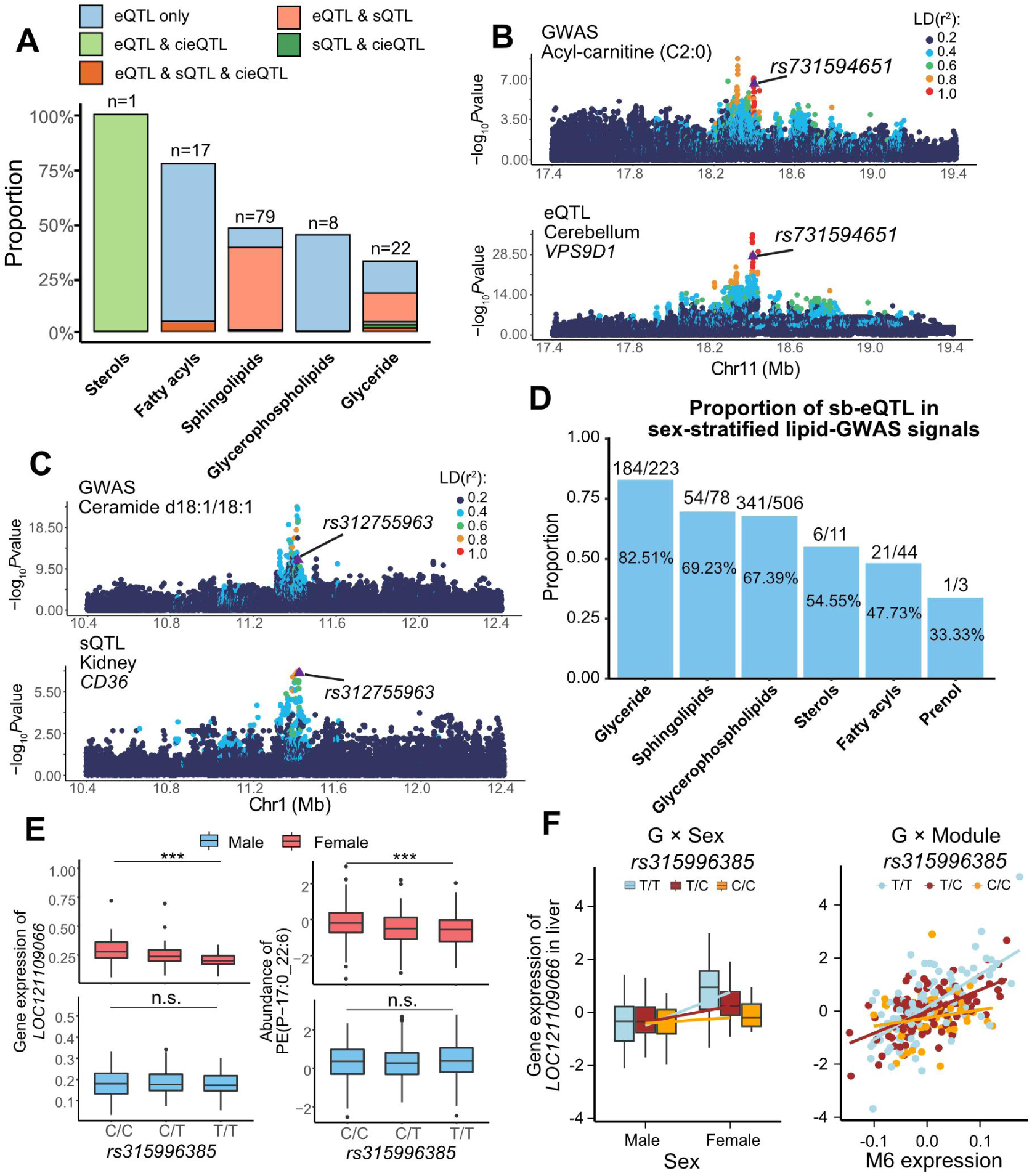
The tissue- and sex-specific gene regulation of serum lipid contents in chickens. **(A)** Proportion of serum lipid GWAS loci (encompassing 865 lipid metabolites) that colocalized with different sets of molQTL, defined by posterior probability of colocalization (PP.H4) > 0.8 using FastENLOC (*120*). **(B)** Colocalization of a serum acyl-carnitine (C2:0) GWAS locus (*rs731594651*) with a *VPS9D1* eQTL in the cerebellum. **(C)** Colocalization of a serum ceramide (d18:1/18:1) GWAS locus (*rs312755963*) with a *CD36* eQTL in the kidney. **(D)** Proportion of sb-eQTL overlapping with sex-stratified serum lipid GWAS signals, with significance (FDR < 0.05) determined via *z*-score test of sex-stratified GWAS results. **(E)** Sex-biased eQTL-mediated regulation of serum phosphatidylethanolamine (PE, P-17:0_22:6) levels. A female-specific sb-eQTL (*rs315996385*) regulating *LOC121109066* expression in the liver coincides with a locus associated with female-specific enrichment of PE (P-17:0_22:6) in the liver. *** denotes statistical significance at *P* < 0.001. **(F)** The sb-eQTL (*rs315996385*) regulating *LOC121109066* expression in a female specific manner and specifically mediated by a co-expression module, M6, in the liver.

Finally, we explored whether the sex-biased regulatory variants (sb-eQTL) identified above could explain the sexual dimorphism in metabolic traits. We found that 49.22% (221) of sb-eQTL exhibited sex-specific associations with at least one serum lipid. These effects were most prominent in glyceride metabolism, where 82.5% of sb-eQTL showed sexually dimorphic impacts, compared to only 33.3% for prenols (**Fig. 8D and data S13**). A compelling example of this sex-specific metabolic architecture is *rs315996385*, which exhibits a female-specific association with phosphatidylethanolamine (PE 17:0_22:6) levels. This variant also functions as a female-specific sb-eQTL for *LOC121109066* in the liver (**Fig. 8E**). Notably, mediation analysis revealed that this female-specific regulatory effect was significantly driven by a liver-specific co-expression module, which was itself characterized by a strong sex bias (**Fig. 8F**). These results demonstrate that sexual dimorphism in the lipidome is not merely a consequence of hormonal differences, but is partially rooted in a shared genetic architecture where sex-biased variants act through coordinated transcriptional networks and tissue-specific regulatory programs to drive divergent metabolic outcomes between sexes.

### Regulatory atlas elucidates the genetic architecture of complex traits

To demonstrate the utility of the ChickenSexGTEx resource in resolving the molecular basis of complex traits, we conducted GWAS of 26 complex traits in 1,575 chickens (768 males and 807 females). These traits span six diverse functional categories, including 11 behavior, 8 growth, 3 metabolism, 2 reproduction, 1 cardiovascular, and 1 immune performance (**table S3**). Our analysis identified 2,896 trait-associated loci, among which sb-eQTL were significantly enriched for genetic signals underlying rooster comb height—a definitive secondary sexual characteristic—as well as primary growth-related traits, including 45-day body weight and average daily gain (**fig S44A and B**). Colocalization analysis revealed that 12.5 to 36.6% of GWAS signals across all 26 traits were colocalized with at least one molQTL (PP.H4 > 0.8) (**Fig. 9A**). Notably, 77.4% of these colocalized signals were associated exclusively with either eQTL or sQTL. For example, the 90-day body weight locus *rs312346400* colocalized with a vascular eQTL of *RAD51B*, whereas another weight-associated variant colocalized solely with a vascular sQTL of *SPIRE1* (**fig 44C and D**). In rare cases of pleiotropic regulation, such as the feed conversion ratio (FCR) locus *rs738482813*, we identified a variant that colocalized with both eQTL and sQTL of *ST3GAL4* across multiple intestinal segments (**Fig. 9B and fig. S45A**), identifying it as a promising candidate for chicken growth efficiency (*70*). Beyond bulk tissue analysis, the integration of cieQTL provided an additional layer of resolution, colocalizing with 369 additional GWAS loci that were entirely independent of bulk eQTL signals (**Fig. 9C**). For example, a GWAS locus of adipose weight on chromosome 4 colocalized strictly with a cieQTL (radial glial cell) rather than bulk eQTL of *TRAF3IP1* in the cerebellum (**Fig. 9D**).

**Fig. 9.**
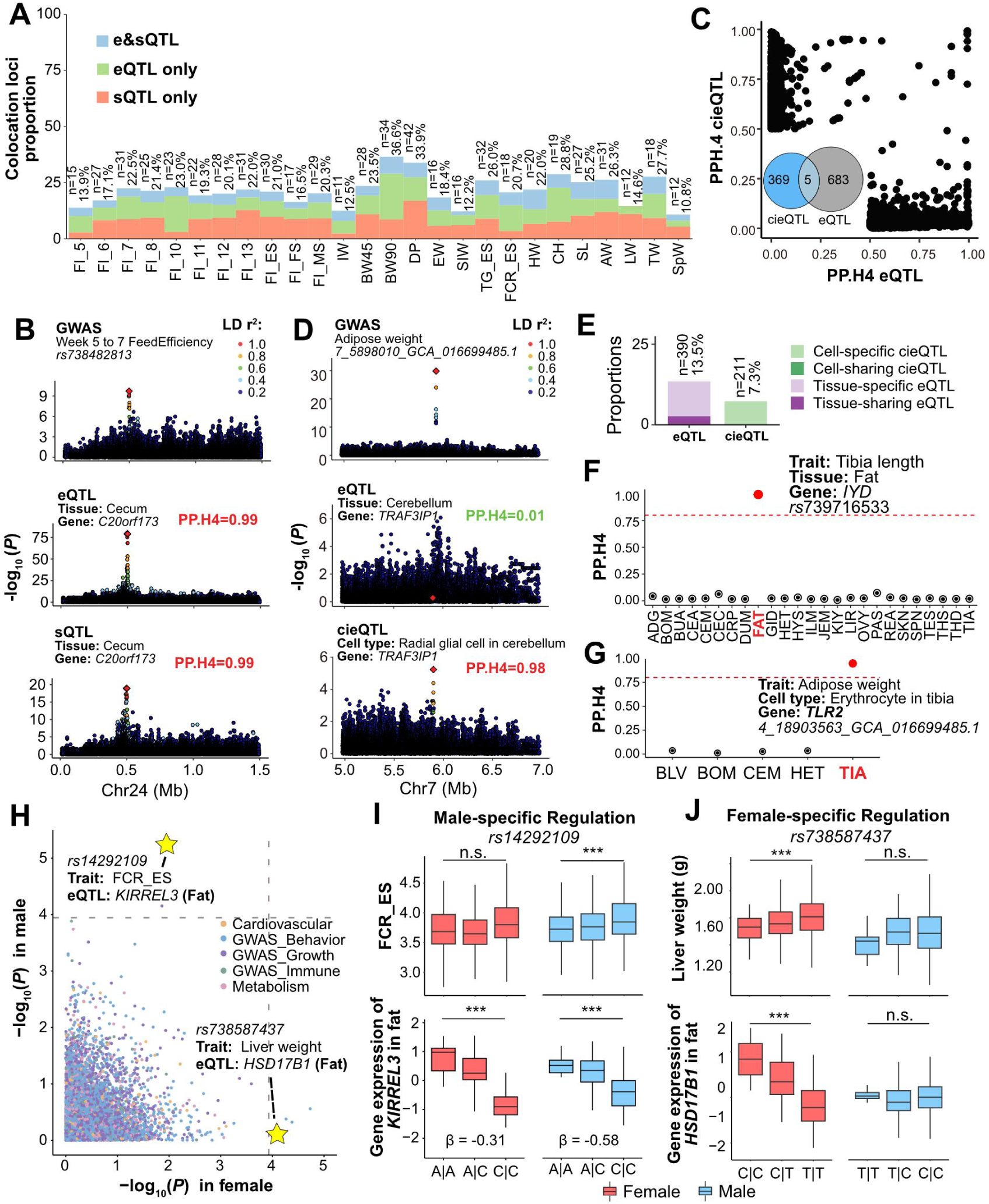
Context-dependent genetic regulation of complex traits. **(A)** Proportion of GWAS loci that colocalized with molQTL (e&sQTL, eQTL only, and sQTL only) across 26 complex traits, defined by posterior probability of colocalization PP.H4 > 0.8. **(B)** Colocalization of GWAS locus (*rs738482813*) for week5 to 7 feed efficiency with a pleiotropic variant that regulates both the expression and splicing of *C20orf173* in the cecum. **(C)** Summary of GWAS loci colocalized by bulk eQTL and cell-type-interaction (cieQTL). Inserted venn plot shows that only 1.34% (5/374) of cieQTL colocalized signals are detected by the bulk eQTL. **(D)** The GWAS locus, 7_5898010_GCA_016699485.1 on chromosome 7, associated with chicken adipose weight significantly (PP.H4 = 0.98) colocalized with the cieQTL of *TRAF3IP1* in radial glial cell but shows no colocalization with the bulk eQTL of *TRAF3IP1* in the cerebellum. **(E)** Proportion of GWAS loci colocalized with tissue/cell-specific or -shared eQTL (cieQTL). **(F)** Tissue-specific eQTL colocalization example. A GWAS locus *rs739716533* of Tibia length shows colocalization with the eQTL of *IYD* in the fat exclusively. The red dash line indicates PP.H4 = 0.8. **(G)** Cell-type-specific cieQTL colocalization example. A GWAS locus of adipose weight, *4_18903563_GCA_016699485.1*, shows colocalization with the erythrocyte *TLR2* cieQTL in tibia tissue exclusively. The red dash line indicates PP.H4 = 0.8. **(H)** Comparison of association tests (-log_10_*P*) between all 449 sb-eQTL and 26 complex traits in male- and female-stratified analysis. Dash line indicates the significance cutoff set as 1.11×10^−4^ (0.05/449) for both sex-stratified association tests. **(I)** Example of male-specific genetic controls underlying complex traits. Genetic variant *rs14292109* shows the male-specific association with week 5-7 feed efficiency, which also exhibits a larger effect size in males compared to females in regulating *KIRREL3* gene expression in fat tissue. **(J)** Example of female-specific genetic controls underlying complex traits. Genetic variant *rs738587437* shows the female-specific association with chicken liver weight, which also exhibits a sex-specific regulation of the estrogen activation gene *HSD17B1* expression in fat tissue.

GWAS-colocalized eQTL were significantly more likely to be tissue-specific (10.81%) than widely shared across tissues (2.66%) (**Fig. 9E**). Vascular and renal tissues showed the largest number of tissue-specific colocalizations (**fig. S45B**). Of note, 35.63% of the GWAS loci for FCR were colocalized with intestinal molQTL (**fig. S44A**), highlighting the importance of the intestine in nutrient digestion and absorption. This includes *rs3386551955*, which was associated with *CLCA1* expression exclusively in the jejunum (**fig. S45C**), consistent with the known role of *CLCA1* in maintaining mucus barrier homeostasis and optimizing nutrient absorption (*71, 72*). Furthermore, we identified a adipose-specific eQTL of *IYD* associated with tibia length (**Fig. 9F**), providing a molecular basis for the known endocrine crosstalk between adipose tissue and bone development (*73–77*). Regarding tissue-shared eQTL, a feed efficiency signal *rs738482813* on chromosome 24 was colocalized with the eQTL of *KIRREL3* in multiple intestinal and metabolic tissues. This robust association was further validated by a transcriptome-wide association study (TWAS) (**fig. S45D and E**). Cell-type-specific interactions further refined these mechanisms. GWAS loci were 70-fold more likely to colocalize with cell-specific cieQTL than with cell-shared signals (7.18% vs. 0.10%) (**Fig. 9E**). For instance, an adipose weight locus (*4_18903563_GCA_016699485.1*) colocalized solely with a cieQTL of *TLR2* in tibial erythrocytes, but not in the erythrocytes of other tissues (**Fig. 9G and fig. S45F**). It has long been known that host immunity highly determines the balance of fat metabolism (*78, 79*). Given that *TLR2* triggers NF-κB signaling to modulate hematopoiesis and lipid oxidation (*80, 81*), his localized immune-metabolic axis underscores the role of the bone marrow niche in regulating systemic lipid metabolic balance.

Finally, we explored whether sex-biased gene regulation serves as the molecular driver of sexual dimorphism in complex traits. By integrating 449 sb-eQTL with sex-stratified GWAS of 26 complex traits, we identified six variants where sex-biased regulatory effects translated into sex-specific phenotypic outcomes (sex-stratified association *P* < 1.11×10^−4^; **Fig. 9H**). For instance, both TWAS and bulk eQTL colocalization identified that the expression of *KIRREL3* was negatively associated with the feed efficiency (**fig. S45D and E**). However, the sb-eQTL *rs14292109* showed the male-specific association with feed efficiency, likely driven by the heightened inhibitory effect size observed in males (*β* = −0.58) relative to females (*β* = −0.31) (**Fig. 9I**). A striking example of female-specific genetic architecture was observed at the *HSD17B1* locus. The female-specific sb-eQTL *rs738587437*, which regulates *HSD17B1* expression in adipose tissue, showed a corresponding female-specific association with liver weight (**Fig. 9J**). As a pivotal regulator of steroidogenesis, *HSD17B1* activates estrogens while inactivating androgens (*82, 83*). Given that estrogen deficiency is a hallmark of hepatic fat over-accumulation, particularly in post-menopausal women and avian models, we hypothesized a conserved mechanism in chickens (*83–85*). More specifically, the T allele of *rs738587437* may exert a sex-specific inhibitory effect on *HSD17B1* expression, blunting local estrogen activation and subsequently triggering lipid over-deposition, manifested as increased liver weight exclusively in females.

### Conserved regulatory syntax links chicken molQTL to human complex traits

To evaluate the translational relevance of the ChickenSexGTEx atlas and quantify the evolutionary constraints on regulatory genome, we systematically compared chicken molQTL with human functional genomic and phenotypic datasets (**Fig. 10A**). Using UCSC LiftOver (*86*), we mapped 175,807 and 83,556 fine-mapped eQTL and sQTL from 32 chicken tissues onto the human reference genome (hg38), yielding 75,143 and 38,276 orthologous variants between humans and chickens. Both eQTL and sQTL were evolutionarily constraint at significantly higher rates than MAF-matched random variants (**Fig. 10B**), and sQTL exhibited a significantly higher liftover rate than eQTL (11.37% ± 0.98% vs. 9.56% ± 0.46%; *P* = 1.23×10^−13^).

**Fig. 10.**
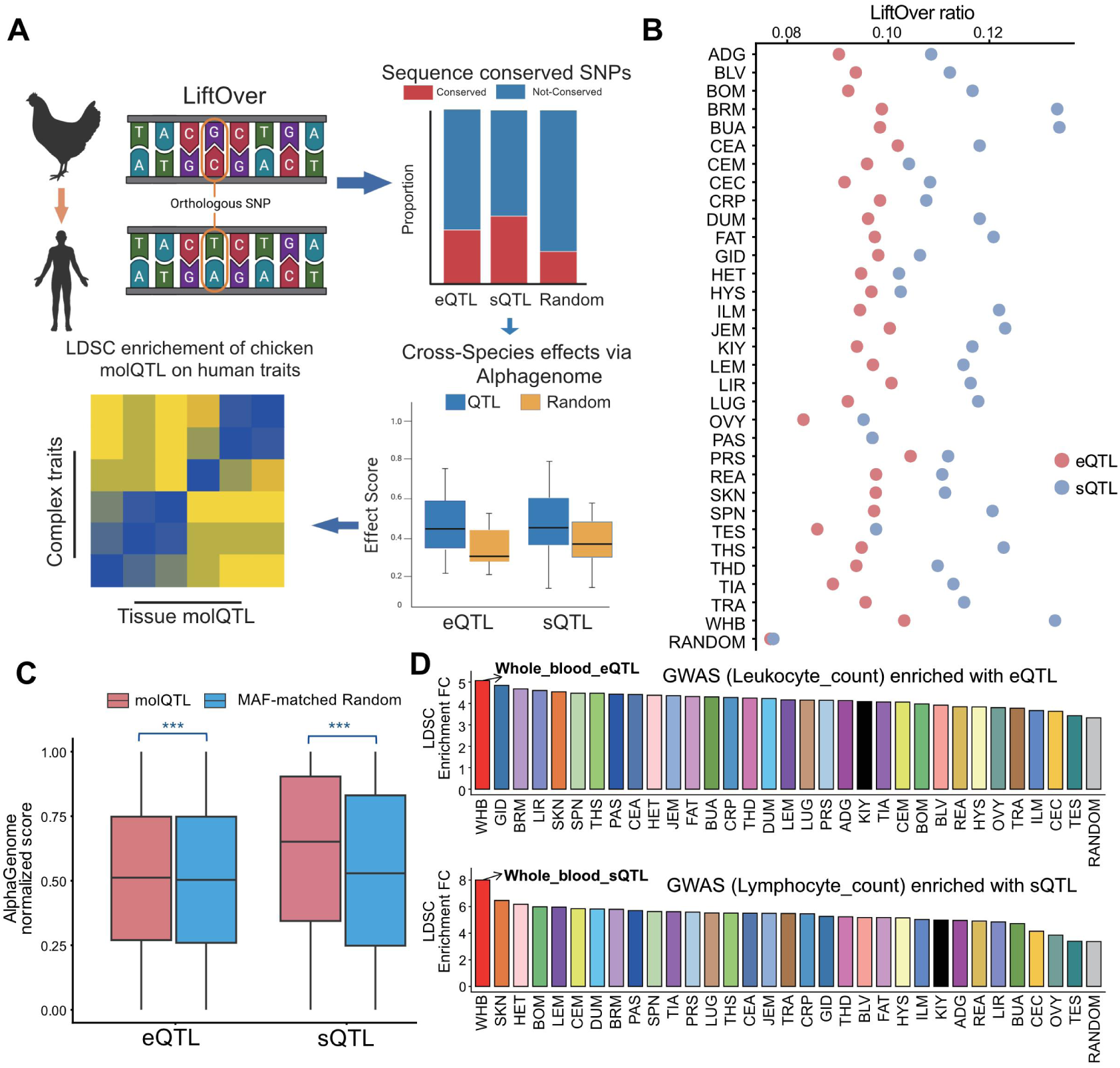
Evolutionary conservation analysis of genetic regulatory variants between chickens and humans. **(A)** The workflow of the evolutionary conservation analysis. **(B)** LiftOver ratio of fine-mapped eQTL and sQTL identified across 32 tissues in chickens onto the human genome. The liftOvered SNPs were considered as orthologous sites between chickens and humans. **(C)** Comparison of AlphaGenome normalized scores between human orthologous sites of chicken e/sQTL and minor allele frequency (MAF) matched random loci. *** represents the significance of *P* value < 0.001. **(D)** The heritability enrichment (calculated by S-LDSC) of orthologous tissue-specific regulatory variants (top panel, eQTL; bottom panel, sQTL) for leukocyte counts in humans.

We next investigated whether these evolutionarily conserved sites maintain their functional “syntax” in the human genome. Using AlphaGenome (*87*), we predicted the impact of chicken molQTL orthologs on human genome function including gene expression and epigenetic modifications. The orthologs of chicken eQTL showed significantly higher predicted impact scores on human gene expression profiles than MAF-matched random controls (*P* = 3.97×10^−18^). This functional conservation was even more pronounced for splicing; where the impact scores on human splicing profiles significantly exceeded those of random controls (*P* = 7.36×10^−131^) (**Fig. 10C**). These data indicate that despite the ∼310 million years of divergence between avian and mammals, the DNA sequence features governing splicing and expression regulation might be conserved across vertebrates.

Finally, using the Stratified Linkage Disequilibrium Score Regression (S-LDSC) (*88, 89*), we explored whether heritability of 133 human complex traits is enriched in the human orthologous variants of chicken molQTL. Orthologs of chicken regulatory variants exhibited a tissue-specific enrichment for human trait heritability (**Data S14**). For instance, chicken whole blood eQTL showed the highest enrichment for human white blood cell leukocyte count, while chicken thymus eQTL specifically pinpointed human basophil percentage. Intriguingly, sQTL in chicken leg muscle were highly enriched for the heritability of human waist-to-hip ratio, indicative of a conserved splicing logic governing metabolic fat distribution (**Fig. 10D**). Collectively, these findings demonstrate that the tissue-specific regulatory networks in chickens recapitulate the molecular basis of human complex traits. This cross-species alignment not only validates the chicken as a potential model for functional genomics but also suggests that avian regulatory maps can be used to pinpoint causal variants in the human genome.

## Discussions

In summary, by integrating 280 WGS with nearly 8,000 bulk RNA-seq and 93 sc/nRNA-seq of 32 primary tissues, we established the ChickenSexGTEx resource as a high-resolution atlas of sexual dimorphism in chickens, resolving the complex interplay between genetic variation, cellular heterogeneity, and tissue-specific gene expression. While landmark efforts like the Human GTEx project or the ChickenGTEx pilot have fundamentally reshaped our understanding of regulatory genome (*25, 90*), they are inherently constrained by postmortem tissue degradation or uncharacterized environmental noise. By contrast, our framework leveraged a rigorously controlled experimental design, rearing a local chicken population under uniform conditions and performing tissue collection within a strict 20-minute window. This minimized environmental confounding to an unprecedented degree. A fundamental challenge in transcriptomics is distinguishing between regulatory variation and shifts in cellular populations. Our cell type deconvolution demonstrated that “compositional bias“—such as female-biased plasma cells versus male-biased macrophages—accounts for a significant portion of apparent transcriptomic dimorphism. By accounting for cell-type proportions, we mapped 10,937 cieQTL that were undetected in bulk data. While cell-type deconvolution is a cost-effective tool for large-scale studies, it is fundamentally an inference-based approach that might miss the biological nuance that only single-cell resolution can capture. For instance, we only considered major cell types, as the cell type deconvolution struggles to estimate rare cell populations which might be very essential for complex traits.

Although many genes were differentially expressed between sexes, we only uncovered 340 genes governed by 449 sex-specific regulatory variants, consistent with findings in human GTEx (*22, 91*). Our mediation analysis revealed that ∼80% of these sex-specific effects are orchestrated by Z-linked transcription factors, co-expression networks, and cell type proportions, suggesting a hierarchical model where sex chromosomal dosage and gene interactive signaling converge to fine-tune specific regulatory loci (*14, 92*). Moving beyond SNPs, we demonstrated that SVs are pivotal, yet frequently overlooked, modulators of the transcriptome. Compared to SNPs, SVs exhibited larger regulatory effect size and higher tissue specificity, with sex-biased SV-eQTL concentrated in immune and endocrine tissues. These findings suggested the potential larger impact of SVs in regulating gene expression, thus purifying selection will work stronger against SVs for general expressed genes and therefore be more tissue specific (*93, 94*).

While ChickenSexGTEx provides a foundational atlas, several limitations remain. Our reliance on short-read WGS restricts our ability to detect complex, large-scale structural variants and repetitive elements, which likely harbor additional regulatory signals accessible only via long-read sequencing technologies (e.g., ONT and PacBio). Similarly, the use of short-read RNA-seq limited our capacity to explore the genetic regulation of individual isoforms, which may exhibit distinct regulatory architectures from our gene-level eQTL. Furthermore, our data represents a physiological snapshot at the age of 90 days. Because gene regulation is inherently dynamic, we likely miss transient events critical for embryonic development and pubertal maturation (*95–97*).

While our fine-mapping and colocalization analyses provide statistical evidence for causal variants of complex traits, we lack high-throughput experimental validation—such as CRISPR-Cas9 editing or reporter assays—to definitively confirm the molecular mechanisms of the identified loci (*98–101*). Despite these caveats, ChickenSexGTEx provides a foundational atlas for decoding the molecular logic of sexual dimorphism, offering profound insights into how genetic variation, cellular context, and sex interact to drive vertebrate phenotypic diversity.

## Materials and methods summary

Detailed methods are in Supplementary Materials. Briefly, this study employed a local chicken population maintained for eight generations by random mating to limit inbreeding and preserve genetic diversity. A total of 1,400 chicks from the 8th generation were reared to 90 days of age under strictly controlled conditions, including phase-specific housing, temperature, lighting, and *ad libitum* feeding. From this cohort, 280 (150 males and 130 females) genetically diverse individuals (mean pairwise IBD = 0.018) representing the founder population structure were selected for the ChickenSexGTEx project. In total, 8,263 high-quality samples across 32 tissues, were collected and stored at −80 °C. Another 2,000 chicken cohort were reared under the same conditions to construct the population of subsequent lipid metabolite and complex trait GWAS. All animal procedures were approved by the Institutional Animal Care and Use Committee of Hunan Agricultural University (No. 2025DKJQ020).

Total tissue bulk RNA was extracted using miRNeasy Mini kits with DNase digestion; RNA integrity (RIN ≥8.0) was verified using Agilent TapeStation. Genomic DNA was isolated from whole blood using DNeasy Blood & Tissue kits. Single-cell suspensions were prepared via tissue-specific enzymatic digestion, filtered, and cryopreserved with >95% viability for 10x Genomics library construction. Whole-genome sequencing was performed on 280 samples. SNPs were called jointly and filtered for missingness, minor allele frequency (≥0.05), and quality metrics; phasing was performed using Beagle (*102*). Pangenome graphs were constructed for SV genotyping via vg Giraffe (*103*), BayesTyper (*104*), GraphTyper2 (*105*), and PanGenie (*106*). RNA-seq was conducted for 8,265 tissue samples. Reads were trimmed with Trimmomatic (*107*), aligned using STAR (*108*), and rigorously filtered for mapping quality and gene expression. Sample identity was validated by comparing RNA-derived SNPs with whole-genome sequencing data. Gene expression was quantified as raw counts and TPM; alternative splicing was quantified using LeafCutter (*109*).

Molecular QTL (eQTL and sQTL) mapping was conducted across 32 tissues using linear mixed models implemented in OmiGA (*110*), adjusting for genetic relatedness, population structure, sex, and technical covariates. Independent signals and credible causal variants were identified via stepwise regression and SuSiE (*28*) fine-mapping. Tissue sharing of regulatory effects was analyzed using MashR (*29*). A single-cell transcriptomic atlas was generated across 31 tissues using 10x Genomics, with quality control, clustering, batch correction (Harmony) (*111*), and cell-type annotation using canonical markers and SingleR (*112*), defining 127 distinct cell types. Bulk-tissue cell-type deconvolution was performed using CIBERSORT (*113*) with a custom signature matrix, validated by simulated pseudo-bulk data. Cell-type-interacted eQTL (cieQTL) mapping was conducted using genotype-by-cell-fraction interaction models. Sex-biased gene expression was quantified across 30 non-gonadal tissues using edgeR (*114*) and limma (*115*). Sex-biased modules were identified by WGCNA (*116*); sex-differential cell-type abundance was assessed via t-tests. Sex-biased eQTL (sb-eQTL) were mapped using genotype-by-sex interaction linear models in tensorQTL (*117*). Mediation analysis evaluated whether sex differences in cell-type proportion, co-expression modules, or transcription-factor expression mediated sb-eQTL effects. Serum lipidomic profiling was performed in 1,000 chickens using LC-MS/MS with MRM detection. GWAS for serum lipids and 26 complex traits was conducted using a linear mixed model by GEMMA (*118*). Colocalization between molQTLs and GWAS signals was tested using Bayesian coloc methods. Cross-species conservation of chicken regulatory variants was assessed by liftOver (*86*) to the human hg38 assembly, AlphaGenome (*87*) functional prediction, and LD-score regression (*88*) against human complex-trait heritability.

## ACKNOWLEDGMENTS

We thank all technical and scientific members of the ChickenSexGTEx and Hunan Agricultural University, who participated sample collection and made the data available. **Funding:** This work was supported by the National Key Research and Development Program of China (2024YFF1000100), Hunan Poultry Industry Technology System (HARS-06), National Broiler Experimental Station of the Modern Agricultural Industrial Technology System (CARS-41-Z08); Yuelu Mountain Seed Industry Special Program (YLS-2025-ZY04050), Agricultural Science and Technology Innovation Program (ASTIP-IAS-04-2), National Key Research and Development Program of China (2023YFF1001000), National Natural Science Foundation of China (32402720), and Jiangxi Provincial Natural Science Foundation (20242BAB20307). **Author contributions:** H.Z.: Writing – Original Draft, Methodology, Investigation. B.L.: Data Analysis, Visualization. Y.X.: Investigation and Data Analysis. D.Z.: Data Analysis, Visualization. C.P.: Data Analysis, Visualization. J.T. and H.L.: Investigation and Data Analysis. Q.O., B.L., C.F., Z.S., and X.M.: Investigation. D.G., B.A., J.T., M.G., L.C., H.L., W.Z., X.Z., and B.L.: Methodology. Y.W., J.L., M.W., and X.L.: Web application. D.W., M.G.P., O.M., R.X., Y.I.L., I.I.-L., Y.Y., X.W., D.H., Q.Z., S.G., Z.S., K.X., G.Y., Z.W., Z.L., C.Z., Y.W., C.S., C.D., S.-H.H., N.Y., X.H., H.Z., and X.Z.: Writing – Review & Editing, Data Interpretation. S.S. and G.Z.: Investigation, Writing – Review & Editing, Funding Acquisition. X.H. and L.L.: Conceptualization, Data Analysis, Writing – Review & Editing. P.Z.: Conceptualization, Data Analysis, Supervision, Writing – Original Draft, Writing – Review & Editing. L.F. and X.H.: Conceptualization, Project Administration, Writing – Original Draft, Writing – Review & Editing. **Competing interests:** All authors declare no competing interests. **Data, code, and materials availability:** The deposited number for the raw sequencing data of whole genome sequence, RNA-seq, and sc/snRNA-seq reported in this paper is PRJCA045295. All codes are accessible in github: https://github.com/PengjuZ/Sex-biased-ChickenGtex. All GWAS and molQTL summary statistics results were available to download at https://sexchickengtex.farmgtex.org/download

## Supplementary Materials

Materials and Methods

Supplementary Text

Figs. S1 to S45

Tables S1 to S3

Data S1 to S15

